# Effect of NaCl-Stressed Bt Cotton on the Feeding Behaviors and Nutritional Parameters of *Helicoverpa armigera*

**DOI:** 10.1101/365213

**Authors:** Jun-Yu Luo, Shuai Zhang, Xiang-Zhen Zhu, Ji-Chao Ji, Kai-Xin Zhang, Chun-Yi Wang, Li-Juan Zhang, Li Wang, Jin-Jie Cui

## Abstract

Saline-alkali soil is an arable land resource on which transgenic Bt cotton has been planted on a large scale in accordance with food security strategies, but there are concerns about the insecticidal effects of Bt cotton on target insect pests. In this study, a *Bacillus thuringiensis* (Bt) cotton variety, GK19, and its nontransgenic parent variety, Simian-3, were used as experimental materials to study the effect of the expression of exogenous insecticidal proteins in Bt cotton under NaCl stress on the feeding behavior and nutritional parameters of *Helicoverpa armigera*. The results showed that the expression of exogenous insecticidal proteins in GK19 was significantly inhibited under NaCl stress. However, on GK19 Bt cotton, the feeding, crawling, resting and spinning down of the 5^th^ instar *H. armigera* larvae, as well as the food consumption and feces amount of these larvae, did not markedly differ under different NaCl concentrations. In contrast, the mean relative growth rate (MRGR), relative growth rate (RGR), approximate digestibility (AD), efficiency of conversion of ingested food (ECI) and efficiency of conversion of digested food (ECD) of the larvae decreased markedly in response to NaCl stress. Under the same concentration of NaCl, the nutritional parameters of the bollworm larvae on GK19 Bt cotton or Simian-3 nontransgenic cotton were different. However, the interaction between salt stress and cotton variety had no significant effect on the feeding behavior or nutritional parameters of *H. armigera* larvae. These results may provide a scientific basis for determining the effect of exogenous insecticidal protein expression in Bt cotton under NaCl stress on *H. armigera* and can therefore be useful for the effective application of Bt cotton in saline-alkali soils to prevent and control *H. armigera*.

## Introduction

Soil salinization is an obstacle that cannot be ignored in the sustainable development of agriculture [1–2]. In recent years, due to improper irrigation and poor drainage,approximately 300,000 hectares of cultivated land worldwide has experienced secondary salinization [3]. In China, nearly 37 million hectares of saline-alkali soil, accounting for 4.9% of the nation’s arable land, severely hinder crop production [4]. Cotton is an important economic crop and the pioneer crop for saline-alkali lands. The rational development and utilization of saline-alkali land resources are highly important for agricultural production.

*Bacillus thuringiensis* (Bt) cotton is commercially cultivated on a large scale in many countries [5–7], as it can effectively control specific lepidopteran pests, reduce dependence on chemical insecticides and protect both beneficial arthropod populations and the environment [8–10]. Although no target pests are beyond the control of extensive Bt cotton use [11–12], many reports have indicated that the effects of Bt cotton on lepidopteran pests differ over the cotton growing season [13] and that the Bt insecticidal protein content is correlated with cotton variety [14–15], cotton growth stage [11,16], and plant parts [17–24]. With respect to the response of Bt cotton to abiotic stress, some studies have investigated the effect of salt stress on the insecticidal protein expression in Bt cotton [19, 20,22, 24], but few studies have investigated the effects of salt stress on both the expression of insecticidal proteins in Bt cotton and the nutritional levels of lepidopteran pests.

Under the context that arable land for grain production is diminishing and that cotton must be planted on saline-alkali land, a salt stress test on Bt cotton plants was simulated in the laboratory to study the exogenous insecticidal protein expression in Bt cotton under NaCl stress and the associated interactions with cotton bollworm (*Helicoverpa armigera*). The results provide a scientific basis for the prevention and control of bollworm in Bt cotton fields in saline-alkali soils.

## Materials and methods

### Plants and insects

Bt cotton variety GK19 and its nontransgenic parent variety Simian-3 were provided by the Institute of Plant Protection and Soil Science, Hubei Academy of Agricultural Sciences, China. GK19 and Simian-3 seeds were sown in germination boxes. After both cotyledons of the cotton seedlings were open and spread flat, the cotton roots were washed, and each seedling was transferred to a hydroponic growth box containing nutrient solution. The growth boxes were subsequently placed in a plant culture room (temperature 25±2 °C, humidity 50-60%) until four true leaves grew. Bollworm larvae were provided by the Institute of Cotton Research of the Chinese Academy of Sciences (CAAS). The bollworm larvae were reared with artificial feed [25] in a light incubator under the 60±5% relative humidity, 2±1 °C, and a 14:10 (light:darkness) photoperiod.

### NaCl stress treatment

The NaCl stress treatments consisted of 3 concentration gradients, namely, 0 mmol L-1, 75 mmol L^−1^ and 150 mmol L^−1^, and each test was repeated 3 times. The cotton nutrient solution and NaCl levels were renewed every 2 days after the initial nutrient and stress application. The nutrient solution was prepared, and the cotton plants were cultivated in accordance with the methods of Jiang et al.[19]; tests were performed for 8 days after the NaCl stress treatment.

### Detection of exogenous protein expression

Unfolded leaves were collected from the tops of GK19 Bt cotton plants under different levels of NaCl stress. For each treatment, 5 leaves were collected. The leaves were then mixed evenly, frozen and ground in liquid nitrogen, after which the exogenous insecticidal protein expression was measured using a Cry1Ac/Cry1Ab detection kit (ENVIROLOGIX, USA).

### Survey of bollworm feeding behavior

GK19 Bt cotton and Simian-3 nontransgenic cotton plants under different levels of NaCl stress were collected. For each treatment, 5 plants were collected, and the cotyledonary nodes and roots were cut and discarded from each plant. The remaining plant portions were placed into a plastic cup containing 1% solidified water agar. A single recently hatched bollworm larva was placed onto each plant. Subsequently, the plastic cup was covered with an inverted plastic cup of equal size, and the cups were sealed together with sealing film. After 10 min, when the bollworm had stabilized, it was observed. This test was repeated 3 times; a total of 15 cotton plants were used. The feeding statuses (feeding, crawling, resting and spinning down) of the bollworm larvae under the different treatments were observed and recorded every 30 min for 6h.

### Determination of the nutritional parameters of the bollworms

To determine the nutritional status of the bollworms, 5^th^ instar larvae were used. Leaves were collected from the tops of the GK19 Bt cotton and Simian-3 nontransgenic cotton and then placed into clean, sterilized glass test tubes (15 cm×Φ5 cm). Afterward, similarly sized bollworm larvae were placed onto the cotton leaves, and each test tube was plugged with a cotton swab to prevent the bollworms from escaping. One 5^th^ instar bollworm larva was placed in each tube, and larvae were reared for 2 d; 30 larvae were examined per treatment, and every 10 larvae represented one replicate. Before rearing, both the weight of the bollworm larvae and the fresh weight of the cotton leaves were measured; after rearing, the weight of the bollworm larvae and the dry weight of the remaining cotton leaves were measured after drying at 80 °C for 72 h. In addition, the fresh weight of the feces in the test tubes were measured. Ten leaves without bollworms served as blank controls for each treatment. The following nutritional effect parameters were calculated: the mean relative growth rate (MRGR) [26], relative growth rate (RGR), relative metabolic rate (RMR), relative consumption rate (RCR), efficiency of conversion of ingested food (ECI), efficiency of conversion of digested food (ECD) and approximate digestibility (AD) [27–28]. The food consumption, amount of feces, larval weight and weight gain of the bollworms were measured.

### Data analysis

SPSS 17.0 statistical analysis software for Windows (SPSS, Chicago, IL, USA) was used to analyze significant differences in the sample data. The Bt insecticidal protein expression, nutritional effect parameters and feeding behavior parameters of the bollworms were analyzed via one-way ANOVA with SPSS 17.0. Significant differences between treatments were tested using LSD tests.

## Results

### Effect of NaCl stress on the expression of exogenous insecticidal protein in Bt cotton leaves

With the increase in NaCl concentrations, the expression of exogenous proteins in the Bt cotton leaves tended to decrease (Fig. 1). Compared with 0 mmol L^−1^ NaCl stress, 75 mmol L^−1^ and 150 mmol L^−1^ NaCl stress caused 23.16% (*p*=0.022) and 59.01% (*p*=0.003) lower exogenous protein expression in Bt cotton leaves, respectively. Compared with 75 mmol L^−1^ NaCl stress, the exogenous protein expression in Bt cotton leaves under 150 mmol L^−1^ NaCl stress decreased by 46.66% (*p*<0.001). Together, these results showed that NaCl stress could significantly inhibit the exogenous protein expression in Bt cotton and that the decrease in the exogenous protein expression was related to the degree of salt stress.

**Fig. 1.**
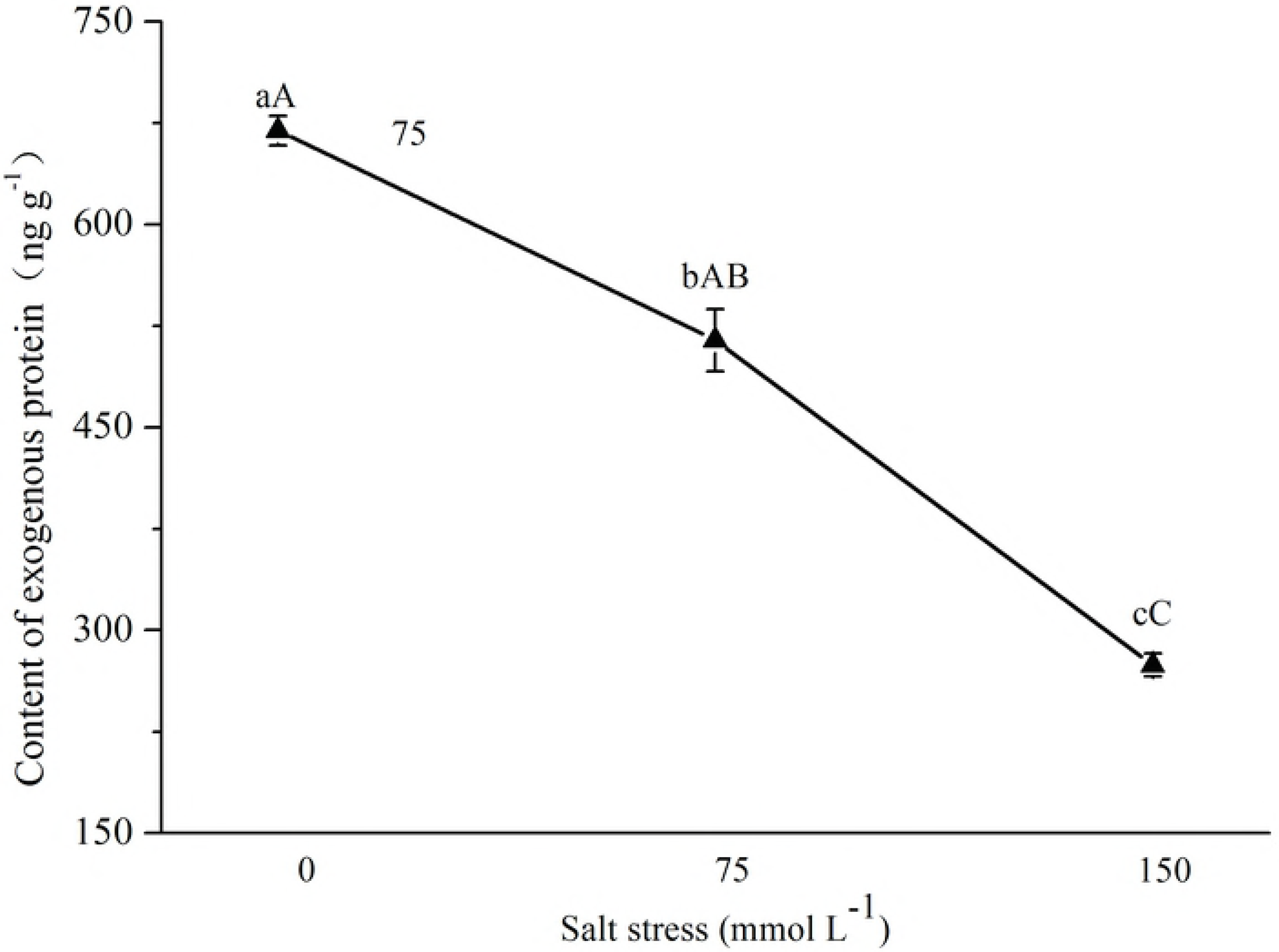
Effect of NaCl stress on exogenous insecticidal protein expression in Bt cotton. Mean (±SD) exogenous protein contents of cotton bollworms (*Helicoverpa armigera*) fed excised transgenic Bt (Bt) cotton plants grown under NaCl stress. Different lowercase letters indicate significant differences between NaCl stress treatments (LSD test: *p*<0.05).

### Effect of NaCl-stressed Bt cotton on the feeding behavior of bollworms

The effect of NaCl-stressed Bt cotton on the feeding behavior of bollworm larvae is shown in Fig. 2. When GK19 Bt cotton and Simian-3 nontransgenic cotton plants were under NaCl stress, the probability that bollworm larvae would feed on them decreased as NaCl concentration increased, but the difference between treatments was not significant. Compared with the feeding probability of bollworm larvae on Simian-3 nontransgenic cotton plants, that on GK19 Bt cotton plants was low overall. Under 0 mmol L^−1^ NaCl stress, the feeding probability of bollworm larvae on GK19 was 42.65% (*p*=0.037) lower than that on Simian-3. Furthermore, the feeding probability of bollworm larvae did not significantly differ under 75 mmol L^−1^ and 150 mmol L^−1^ NaCl stress; the probabilities of crawling, resting and spinning down for larvae on Bt cotton were not significantly different from those for larvae on nontransgenic cotton, even under different levels of NaCl stress. These results show that NaCl stress did not significantly affect the feeding behavior of bollworm larvae on cotton plants.

**Fig. 2.**
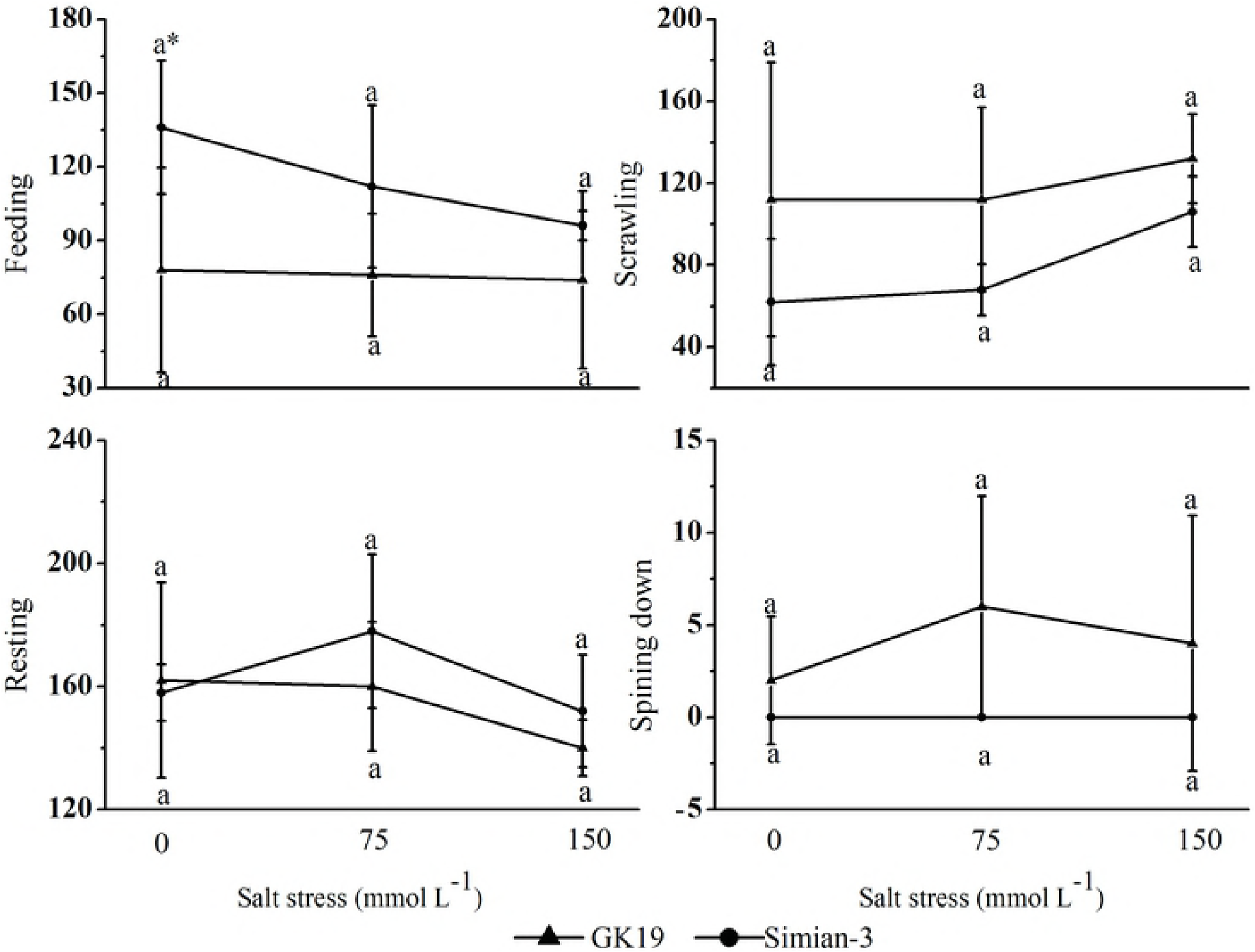
Effect of NaCl-stressed Bt cotton on the feeding behavior of bollworm larvae. (A) Mean (±SD) feeding, (B) crawling, (C) resting and (D) spinning down behavior of cotton bollworms (*Helicoverpa armigera*) fed excised transgenic Bt (Bt) and nontransgenic (Non) cotton plants grown under NaCl stress. Different lowercase letters indicate significant differences between NaCl stress treatments, and * indicates significant differences between the cotton varieties under the same NaCl stress treatment (LSD test: *p*<0.05).

### Effect of NaCl-stressed Bt cotton on the nutritional effect parameters in bollworms

#### Food consumption and the amount of feces

The cotton variety and NaCl stress had marked effects on the food consumption and amount of feces of bollworm larvae (Fig. 3). Under the same NaCl stress condition, the food consumption and amount of feces of bollworm larvae on GK19 Bt cotton were significantly lower than those on Simian No. 3 nontransgenic cotton (*p_0_*=0.000, *p_75_*=0.000, *p_150_*=0.001). Compared to the 0 mmol L^−1^ NaCl treatment, the food consumption and amount of feces of bollworm larvae on the Bt cotton under 75 mmol L-1 and 150 mmol L^−1^ NaCl both first increased but then decreased, and there was no significant difference among the stress treatments. However, the food consumption and amount of feces of the cotton bollworm larvae on nontransgenic cotton were decreased, and under 150 mmol L^−1^ NaCl, these characters decreased by 25.88% and 23.06% (*p*_75_=0.002, *p*_150_=0.021), respectively. Together, these results showed that the introduction of exogenous genes significantly inhibited the food consumption and amount of feces of bollworm larvae on cotton. However, as the NaCl concentration increased, no significant differences in the bollworm food consumption and amount of feces were observed on either cotton variety.

**Fig. 3.**
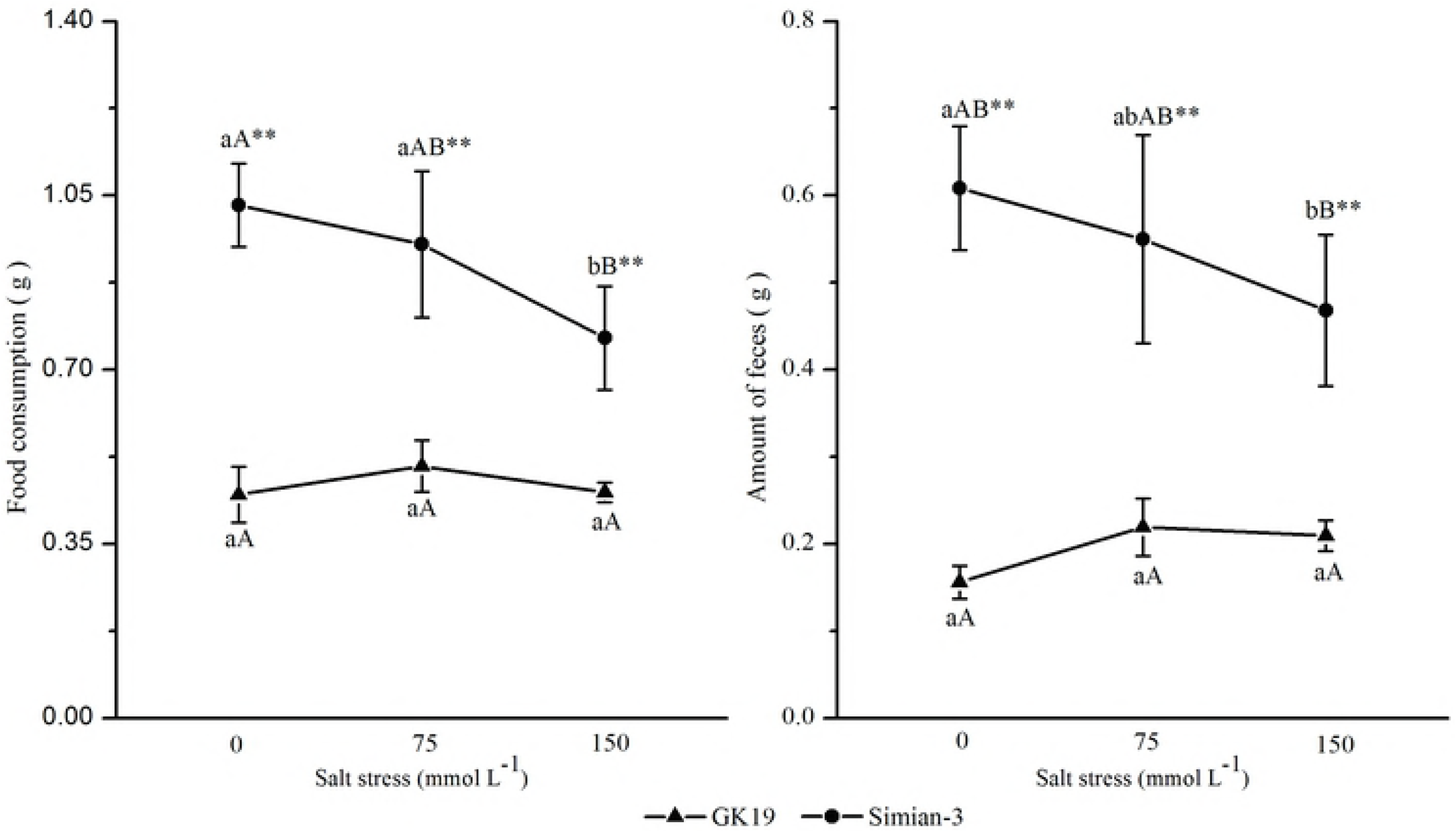
Effect of NaCl-stressed Bt cotton on the food consumption and amount of feces of bollworm larvae. (A) Mean (±SD) food consumption and (B) amount of feces of cotton bollworms (*Helicoverpa armigera*) fed excised transgenic Bt (Bt) and nontransgenic (Non) cotton plants grown under NaCl stress. Different lowercase letters indicate significant differences between NaCl stress treatments, and * or ** indicates significant differences between cotton varieties under the same NaCl stress treatment (LSD test: *p*<0.05 and *p*<0.01, respectively).

#### Nutritional effect parameters

The cotton variety and NaCl stress significantly affected the MRGR, RGR, RCR and relative consumption rate (RMR) of the bollworm larvae (Fig. 4). Under the same NaCl stress level, the MRGR, RGR and RCR of bollworm larvae feeding on GK19 Bt cotton were significantly lower than those of bollworm larvae feeding on Simian-3 nontransgenic cotton (*p*<0.001), but the RMR of the former was significantly higher than that of the latter (*p*>0.05). Compared with the MRGR and RGR under 0 mmol L-1 NaCl stress, those of bollworm larvae feeding on both varieties of cotton under 75 mmol L^−1^ and 150 mmol L^−1^ NaCl stress decreased, but the MRGR (*p*_75_=0.046, *p*_150_=0.014) and RGR (*p*_75_=0.027, *p*_150_=0.050) of bollworm larvae feeding on Bt cotton and nontransgenic cotton under 150 mmol L^−1^ NaCl stress significantly decreased.

**Fig. 4.**
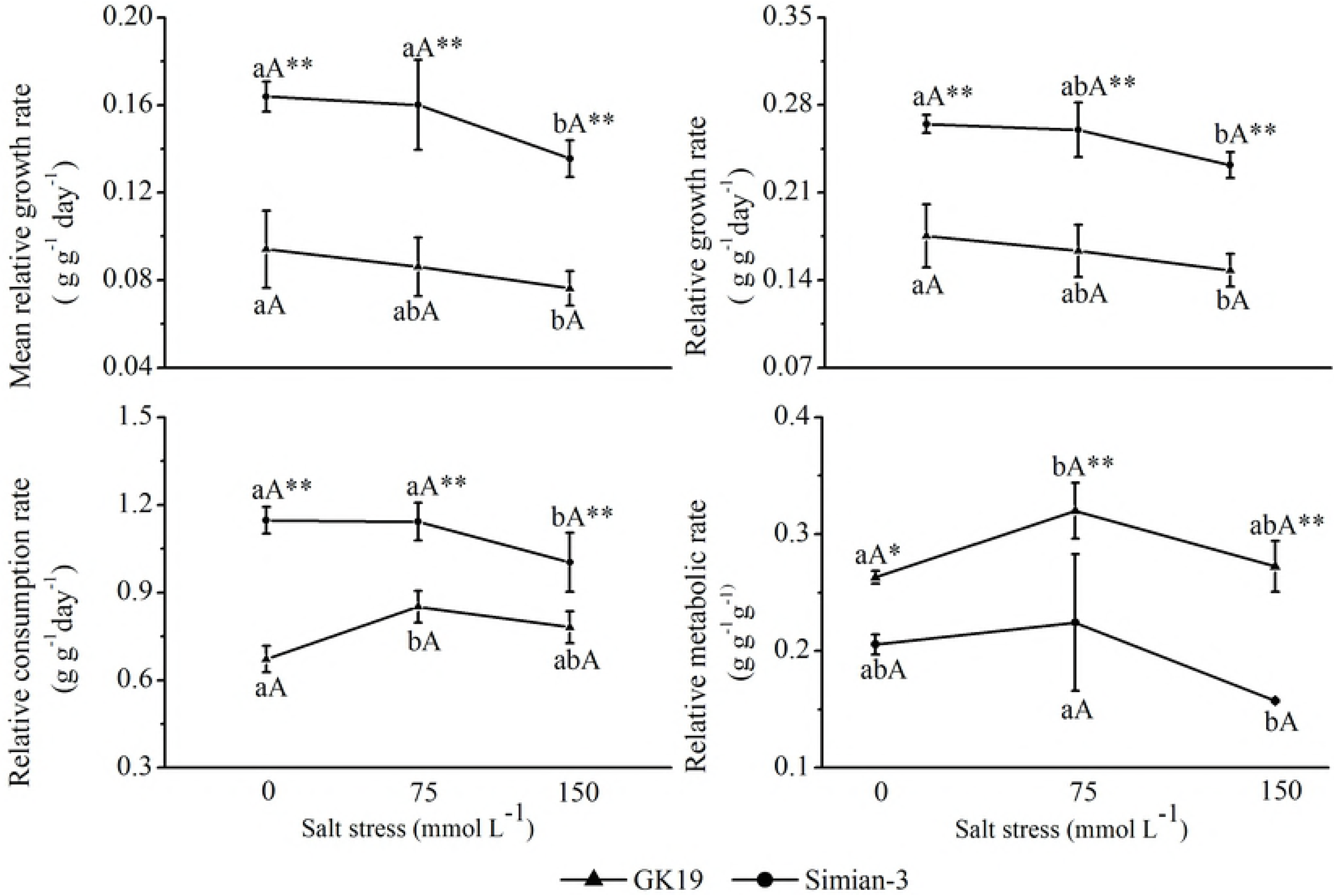
Effect of NaCl-stressed Bt cotton on the MRGR, RGR, RCR and RMR of bollworm larvae. (A) Mean (±SD) relative growth rate (MRGR), (B) relative growth rate (RGR), (C) relative consumption rate (RCR), and (D) relative metabolic rate (RMR) of cotton bollworms (*Helicoverpa armigera*) fed excised transgenic Bt (Bt) and nontransgenic (Non) cotton plants grown under NaCl stress. Different lowercase letters indicate significant differences between NaCl stress treatments, and * or ** indicates significant differences between cotton varieties under the same NaCl stress treatment (LSD test: *p*<0.05 and *p*<0.01, respectively).

The RCR and RMR of bollworms feeding on GK19 Bt cotton significantly differed (*p*_RCR_=0.011, *p*_RMR_=0.027) under 75 mmol L^−1^ NaCl stress. In addition, the RCR of bollworms feeding on Simian No. 3 nontransgenic cotton significantly differed (*p*=0.014) under 150 mmol L^−1^ NaCl stress. Overall, these results showed that both cotton variety and NaCl stress can significantly affect the MRGR, RGR, RCR and RMR of bollworm larvae.

The cotton variety and NaCl stress significantly affected the AD, ECI and ECD of bollworm larvae (Fig. 5). Under the same NaCl stress level, the AD of bollworm larvae feeding on GK19 Bt cotton was higher than that of bollworm larvae feeding on Simian-3 nontransgenic cotton; in addition, the ECD and ECI of the former were lower than those of the latter only under 75 mmol L^−1^ and 150 mmol L^−1^ NaCl stress (*p*<0.001). Compared with the AD and ECI under 0 mmol L^−1^ NaCl stress, the AD (*p*_75_=0.004, *p*_150_=0.000) and ECI (*p*_75_=0.000, *p*_150_=0.000) of bollworm larvae feeding on the Bt cotton under 75 mmol L^−1^ and 150 mmol L^−1^ NaCl stress significantly decreased, while the ECD decreased only under 75 mmol L^−1^ NaCl stress (*p*=0.049). The AD, ECI and ECD of bollworm larvae feeding on Simian-3 nontransgenic cotton did not significantly change. Together, these results showed that both the cotton variety and NaCl stress can significantly affect the AD, ECI and ECD of bollworm larvae.

**Fig. 5.**
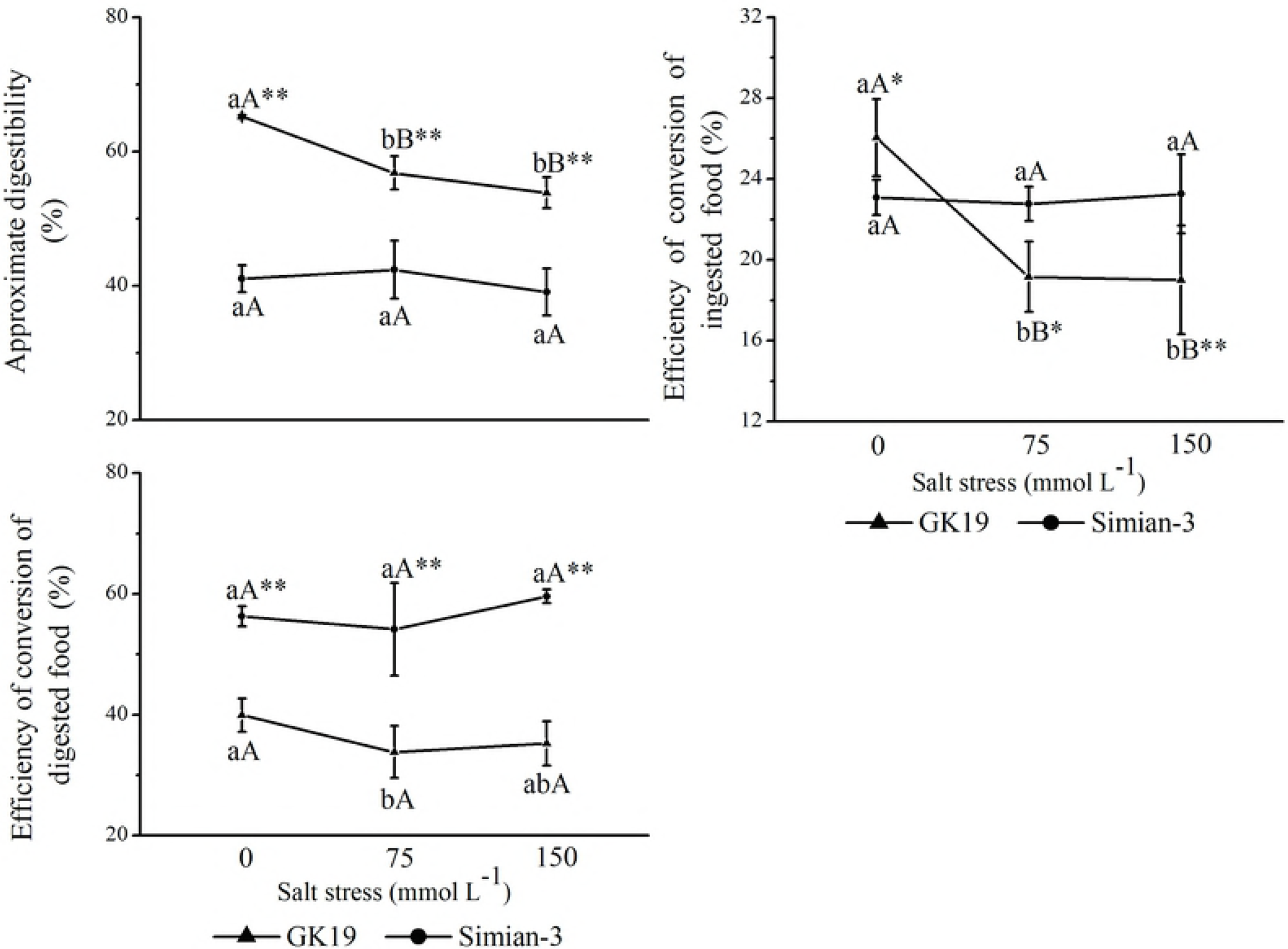
Effect of NaCl-stressed Bt cotton on the AD, ECI and ECD of bollworm larvae. (A) Mean (±SD) approximate digestibility (AD), (B) efficiency of conversion of ingested food (ECI), and (C) efficiency of conversion of digested food (ECD) of cotton bollworms (*Helicoverpa armigera*) fed excised transgenic Bt (Bt) and nontransgenic (Non) cotton plants grown under NaCl stress. Different lowercase letters indicate significant differences between NaCl stress treatments, and * or ** indicates significant differences between cotton varieties under the same NaCl stress treatment (LSD test: *p*<0.05 and *p*<0.01, respectively).

#### Salt stress concentration and cotton variety interaction

Within a certain range, the salt stress concentration, cotton variety, and their interaction have certain effects on the feeding behavior and nutritional parameters of bollworm larvae (Table 1). Salt stress had no significant effect on the probability of bollworm feeding, while cotton variety had a significant effect on the probability of bollworm feeding and crawling; the interaction between salt stress and the cotton variety had no significant effect on the probability of bollworm feeding or crawling. Both the cotton variety and salt stress significantly affected the nutritional parameters of bollworm larvae after feeding, but their interaction had less of an effect on the nutritional parameters. Neither salt stress nor cotton variety had significant interactive effects on the feeding behavior or nutritional parameters of the bollworm larvae.

**Table 1.** Effects of NaCl, cotton variety, and the interaction between NaCl and cotton variety on the feeding behavior and nutritional effect parameters of young cotton bollworm larvae (F-value). * or ** indicates significant differences under the same NaCl stress treatment (LSD test: *p*<0.05 and *p*<0.01).

## Discussion

The insecticidal effect of Bt cotton depends on the expression of exogenous insecticidal proteins, which is correlated with plant variety, plant growth stage, plant tissue and abiotic stress [11,14,16,17,18, 19, 20, 22,24]. In this study, the expression of exogenous insecticidal proteins in the leaves of Bt cotton GK19 was measured under NaCl stress. The results showed that NaCl stress could significantly inhibit the expression of exogenous Bt proteins in the Bt cotton plants, which is consistent with the conclusions of Rao [18], Jiang et al. [19], Luo et al. [20] and Iqbal et al. [22]. Based on their salt stress studies, Li et al. [29] and Luo et al. [24] reported similar conclusions under greenhouse and under natural field conditions. Together, these results show that salt stress can significantly inhibit the expression of exogenous proteins in Bt cotton and that the degree of inhibition is related to the degree and duration of the salt stress.

Different pests exhibit different behavioral responses to transgenic crops; even for the same pest, larvae at different instars respond differently to the same transgenic crop, and the same pest can exhibit different avoidance behaviors to different toxins or genetically modified crop species [30–33]. This study investigated the feeding behavior of newly hatched bollworm larvae on GK19 Bt cotton and its nontransgenic parent Simian-3 under NaCl stress. The results showed that the feeding probability of bollworm larvae on GK19 Bt cotton and Simian-3 nontransgenic cotton under NaCl stress decreased as the NaCl concentration increased; however, there were no significant differences among salt treatments. Compared to the probability of bollworm larvae feeding on Simian-3 nontransgenic cotton, that on GK19 Bt cotton was lower overall and decreased significantly only under no salt stress (0 mmol L^−1^),which is similar to the results under normal planting conditions [31]. However, under salt stress conditions, there was no significant difference in the feeding rate of the newly hatched bollworm larvae on the two varieties of cotton, probably due to the decrease in the bollworm avoidance behavior resulting from the reduced expression of exogenous insecticidal proteins in Bt cotton under NaCl stress [34–36]. The probabilities of bollworm larvae crawling, resting and spinning down on GK19 Bt cotton were not significantly different from those on Simian-3 nontransgenic cotton, and these probabilities did not significantly differ among the NaCl stress treatments. These findings show that NaCl stress had no significant effect on the feeding behavior of bollworm larvae on cotton, as shown by the results of previous experiments on rice brown planthopper (*Nilaparvata lugens*) selection of hosts for oviposition [37].

When grass-feeding insects feed on transgenic insect-resistant plants, food consumption and use may be affected [38]. Whittaker et al. [39] studied the effects of elevated CO_2_ concentrations on plant-herbivore interactions, and their results showed that Bt cotton and elevated CO_2_ concentrations could slow the development of bollworms and consequently reduce the larval weight gain, RGR and MRGR. The present study investigated the effects of GK19 Bt cotton and its nontransgenic parent Simian-3 under NaCl stress on the nutritional parameters of 5^th^ instar bollworm larvae. The results showed that, under the same NaCl stress level, the food consumption and amount of feces of bollworm larvae were significantly lower on GK19 Bt cotton than on Simian-3 nontransgenic cotton, which is consistent with the conclusions of Shobana et al. [40] and Roy et al. [38]. However, there were no significant differences among the stress treatments involving different NaCl concentrations; NaCl stress resulted in significant decreases in the MRGR, RGR, AD, ECI and ECD of the 5^th^instar bollworm larvae, which was essentially consistent with the results of both Whittaker et al. [39] and Chen et al. [41], who investigated the effects of elevated CO_2_ on the nutritional parameters of bollworms feeding on Bt cotton. Somashekara et al. [42] reported that the AD of bollworms decreased after feeding on Bt cotton, while in this study, the AD of 5^th^ instar bollworm larvae feeding on GK19 Bt cotton was much higher than that of 5^th^ instar larvae feeding on nontransgenic cotton, which is consistent with the results that Chen et al. [43] reported in their study about the nutritional parameters of different generations of beet armyworms feeding on transgenic cotton. In addition, the results of the present study suggest that the nutritional parameters of bollworms feeding on transgenic and nontransgenic cotton plants differ under NaCl stress. The variations between these studies and insects may be due to either the response of phytophagous insects to different hosts, which correlates with the growth status of the host plants [44–45,40], or the food compensatory effect of the insects [44]; this question needs to be studied further via both short-term and long-term experiments.

The analysis of the interactive effects revealed that Bt cotton could inhibit the feeding behavior of bollworm larvae; however, salt stress did not affect the chance that larvae would feed on cotton. Although both salt stress and cotton variety significantly affected the nutritional parameters of bollworm larvae after feeding, they had no significant interactive effects on the feeding behavior or nutritional parameters of bollworm larvae. Together, these results are consistent with those reported by Chen et al. [41], who studied the effect of elevated CO_2_ concentrations on the nutritional parameters of bollworms.

Under the background of planting Bt cotton on saline-alkali land, this study investigated the complexity of the response of bollworm larvae to Bt cotton grown under NaCl stress. The results showed that NaCl stress could not induce taxis or reduce the feeding behavior of bollworm larvae on cotton but could affect the growth, development, and nutritional parameters of bollworm larvae. The food utilization rate of phytophagous insects of different species and generations differs highly, and further research is needed to clarify the effect of reductions in the expression of exogenous proteins on bollworms feeding on Bt cotton plants under salt-alkali stress and to quantify the nutritional indicators of herbivores on transgenic insect-resistant plants to determine the impact of these plants on target or nontarget organisms. These findings provide a basis for the control of pests on Bt cotton plants on saline-alkali lands.

## Acknowledgements

We are grateful for the financial support from the National Natural Science Foundation of China (31501253).

